# Enhancement of drought tolerance on guinea grass by dry alginate macrobeads as inoculant of *Bacillus* strains

**DOI:** 10.1101/761056

**Authors:** Jonathan Mendoza-Labrador, Felipe Romero-Perdomo, Juan-Pablo Hernández, Daniel Uribe, Ruth Bonilla Buitrago

**Author notes:** Corresponding Author: Ruth Bonilla Buitrago. This study is in memory of Yoav Bashan (Emeritus Ph.D. in Science). President of The Bashan Foundation. Leader of the Environmental Microbiology Group of Centro de Investigaciones Biológicas del Noroeste (CIBNOR) in México and Founder of Bashan Institute of Science (BIS) Auburn, Alabama, U.S.A.

## Abstract

Drought is responsible for almost 70% of total crop damage in tropical and subtropical countries. In the Colombian Caribbean, drought has caused low availability of Guinea grass (*Megathyrsus maximus*) in forage amount and quality, generating increase in production costs in livestock systems. In this study, we aimed at designing and evaluating dry macro-polymeric inoculants of Bacillus strains and used them successfully to mitigate drought effect on Guinea grass (*M. maximus*). We chose *Bacillus* sp XT13 and *Bacillus megaterium* XT14 strains, previously selected for their capacity to mitigate drought stress on Guinea grass. For the inoculant formulation process, a concentration of 3.2% alginate was used for both microorganisms, selecting the polymer-bacterium ratio of 70:30 for XT13 and 80:20 for XT14. The results of the experiments in greenhouse showed that drought had a negative effect on Guinea grass growth, but the PGPB application with or without macro-polymeric dry matrix improved plant growth significantly. We also observed that co-inoculation of the strains generated a greater beneficial effect than individual inoculation. Interestingly, co-inoculation of XT13 and XT14 under the dry macro-polymeric formulation increased biomass production of the aerial part (7.32%), root biomass, (25.3%), nutritional quality parameters, such as digestibility (3.32%), protein (20.3%); proline accumulation increased (21.06%) and both neutral digestible fiber (2.43%) and APX antioxidant activity (24.2%) decreased with respect to co-inoculation of both bacteria without formulation. These findings have shown that dry macrobeads immobilization has a significantly positive influence on the capacity of XT13 and XT14 bacterial strains to induce drought stress tolerance in Guinea grass.

## 1. Introduction

Climate change has turned out to be a challenge for agriculture and livestock development worldwide due to the increase of biotic and abiotic stress in severity and frequency (Kavamura et al. 2013). Within abiotic stresses, drought affects plant-water relationship, deteriorates growth, influences nutrition and plant development negatively (Schleuning et al. 2016). These effects decrease agriculture cultivation production (Ngumbi and Kloepper 2016) where wheat, rice and barley have been the most reported (Carmen and Roberto 2011). Drought, jointly with climate change, will have caused serious problems for 50% of cultivation land around the world by 2050 (Savo et al. 2016). In the last decade, climate change has made drought seasons lengthy and evident, especially in arid and semi-arid regions (Naveed et al. 2014). While in the dry regions of the Colombian Caribbean, such climate phenomenon has impacted livestock activity negatively with socioeconomic consequences for cattle since productivity of forage quality and quantity based on Guinea grass (*Megathyrsus maximus*) decreases in drought seasons. For example, this situation, affects animal feed, which generates losses in heads of cattle, representing death of 130 thousand bovine specimens in 2013; additionally, losses in milk and meat production were recorded up to 7.6% and 2.2%, respectively, in 2015 (Tapasco et al. 2015).

Based on this problem, different sustainable strategies have been generated to counteract the negative effects of drought, such as developing tolerant varieties and using genetic engineering (Fang and Xiong 2015). Nonetheless, a promising alternative is to induce hydric stress tolerance by using beneficial microorganisms, such as plant growth promoting bacteria (PGPB) (Singh et al. 2016). The bacteria of the genus *Bacillus* have an important role generating physiological and biochemical changes in plants (Yang et al. 2009), providing induced systemic tolerance (IST) to these abiotic stress types (Khan et al. 2019). The mechanisms related with IST include antioxidant defense (APX) (Ghosh et al. 2018; Wang et al. 2012), osmotic adjustment by accumulation of proline (Ghosh et al. 2018), phytohormone production, such as indole-3-acetic acid (AIA) (Spaepen et al. 2014) and defense strategies, such as expressions of genes related to pathogenesis (PR) (Kim et al. 2012). At the same time, reports have demonstrated systemic tolerance induced by PGPB in cultivations under stress by drought, such as cucumber, corn, wheat, bean and chickpea in greenhouse conditions (Sandhya et al. 2010; Wang et al. 2012).

On the other hand, different research studies have asserted that PGPB populations have drastically reduced when inoculated to soil in adverse (drought, salinity and metal toxicity) conditions, losing their biological activity rapidly (Yang et al. 2009). Therefore, providing a protection that allows making plant metabolic activities more efficient is greatly important to stimulate plant growth and generate induced system tolerance. This abiotic stress protection is generated by a polymeric formulation, which is produced through the process of encapsulation with a biodegradable polymer, such as sodium alginate (Bashan et al. 2014). Alginate is currently the most used polymer to immobilize microorganisms (Gonzalez et al. 2018) and several reports have described its beneficial aspects, such as (1) non-toxic and preservative-free; (2) degradation in soil by microbial action; and (3) physical protection from soil predators and environmental stresses by immobilized bacteria in alginate (Bashan et al. 2016; Covarrubias et al. 2012). The immobilization technique is based on a process in which microbial biomass is immobilized due to the polymer reaction with divalent cations (usually Ca ^+2^ in the form of CaC1_2_), forming beads that can be macro- or microbeads (Bashan et al. 2002).

This type of formulations shows beneficial effects for the cells, such as allowing their gradual release to the soil, facilitating their application to the farmer (Singh et al. 2016), generating an adhesive effect on seeds (Bashan et al. 2002; Covarrubias et al. 2012), and creating an adequate microenvironment to preserve their viability and biological activity during lengthy periods of time (Bashan et al. 2014). An inoculant made of alginate macrobeads (2–3 mm) for PGPB was developed over 30 years ago (Bashan 1986), and it has been used for wastewater treatment (de-Bashan et al. 2004), agricultural experimentation worldwide (Bashan et al. 2014; Bashan et al. 2016; Schoebitz et al. 2013), studies of bacteria-microalga interaction (de-Bashan and Bashan 2008; de-Bashan et al. 2016) and bacterial diversity studies in mesquite roots (Galaviz et al. 2018), but it has had limited experimental experience under drought stress so far. Therefore, the objective of this study was to design and evaluate dry macro-polymeric inoculants based on *Bacillus* sp XT13 and *Bacillus megaterium* XT14 strains and use them successfully to increase tolerance to drought effects on Guinea grass (*M. maximus*). The two hypotheses of this research were (1) strains capable of consuming and degrading alginate will have a higher degradation of alginate macrobeads; and (2) Immobilization in macrobeads has a positive influence on the ability of bacteria XT13 and XT14 to induce tolerance to water stress.

## 2. Materials and methods

### 2.1 Microorganisms

*Bacillus* sp XT13 and *B. megaterium* XT14 strains were used in this study for immobilization in alginate macrobeads because they can tolerate drought stress conditions and at the same time promote growth of Guinea grass (*M. maximus*) in hydric stress conditions (Abril et al. 2017; Moreno et al. 2015). These strains were obtained from the Agricultural Microbiology Laboratory strain collection of AGROSAVIA, Colombia.

### 2.2 Growth and cultivation conditions

The strains were reactivated in trypticase soy agar culture medium (TSA, Difco Laboratories, Detroit, MI, USA) at standard conditions: 30 °C for 24 h. Bacterial pre-inoculum was produced aerobically by growing each strain in the log phase at standard conditions subjected to shaking at 150 rpm, cultivating the cells in JM broth (Díaz-García et al. 2015). Then, for spore preparation, we carried out a fermentation process using the pre-inoculum produced in a Rushton (Miniforms, INF-30174, Bottmingen, CH) type turbine-agitated fermenter (capacity 5 L) at 150 rpm, 30 °C with 1 vvm and an effective volume of 3.5 L. At 72 h, a spore concentration of 1.05 × 10^10^ spore mL^−1^ was obtained from each one of the strains that were calculated by making serial dilutions; the count was made using Neubauer chamber (BOECO, DE). Each spore suspension was centrifuged at 4.637 g for 10 min, washed, resuspended in saline solution (NaCl al 0.85% p / v) and stored at 4 °C.

### 2.3 Capacity of degrading alginate and alginolytic activity of *Bacillus* sp XT13 and *Bacillus megaterium* XT14

The capacity of degrading alginate was tested with the methodology proposed by Cruz et al. (2013), utilizing the liquid M.A medium with the following (mg L^−1^) components: NH_4_Cl (50); KH_2_PO_4_ (14); MgSO_4_·3H_2_O (2); CaCl_2_ (4); NaCl (7), supplemented with 10 g L^−1^ of sodium alginate as the only carbon source. The cells of *Bacillus* sp XT13 and *B. megaterium* XT14 strains were extracted by centrifugation 4.637 × g for 10 min, washed three times in saline solution at 0.85%, and their concentration was adjusted to D.O_540 nm_: 0,1 (~10^6^ CFU mL^−1^); then, 20 μl of each one of the adjusted strains were added in 180 μl of M.A. medium. The capacity of the strains to survive was tested in M.A. medium without sodium alginate as the control group.

The experiment was conducted twice and incubated in 96-well microplates at a temperature of 30 ± 1° C for 84 h. The experimental design was composed of six treatments (without (-); with (+) alginate (Alg)): T1: M.A – Alg, T2: M.A + Alg, T3: M.A + Alg + (XT13), T4: M.A + Alg + (XT14), T5: M.A – Alg + (XT13). T6: M.A – Alg + (XT14). Each treatment was performed with 12 replicates. The response variable was the record of bacterial population per hour at OD:540 nm (DR-2000, Hach).

Likewise, an experiment based on viscosity measurement, as an indicator of alginolytic activity, was performed with a Gilmont (Thermo Fisher Scientific, Waltham, MA, USA) viscosimeter using the following equation (1):

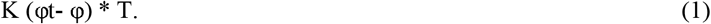

Where density of the stainless-steel ball (φ) is 8.02. The density (φt) associated with the fluid was determined with a pycnometer; K is 35 for tube number 3 used in this test; T is displacement time of the stainless-steel ball in glass tube number 3.

A completely randomized design was used with a factorial arrangement of 2*2. The factors were polymer-bacterium ratio (80:20 and 70:30 vol / vol) and microorganisms (XT13 and XT14) plus a control treatment of sodium alginate at 3.2% without bacteria. The experimental design consisted of five treatments T1: sodium alginate at 3.2%; T2: XT13 strain with a ratio 80:20 (polymer:bacterium), T3: XT13 strain with a ratio 70:30 (polymer:bacterium), T4: XT14 strain with a ratio 80:20 (polymer:bacterium), T5: XT14 strain with a ratio 70:30 (polymer:bacterium) with three replicates each one; 50 mL of each treatment were incubated In 250mL flasks in an orbital shaker at 100 rpm at 28°C during 96 hours’ time. The entire experiment was repeated twice.

### 2.4 Formulation of dry macrobead inoculant

#### 2.4.1 Immobilization

Four prototypes were formulated with the polymer-bacterium (80:20 and 70:30 vol / vol) ratio for each one of the strains. The spore suspensions were obtained in the fermentation process; 200 mL and 300 mL of these suspensions were mixed with 800 mL and 700 mL of sodium alginate suspension (3.2% p/v) for the ratios 80:20 and 70:30, respectively. The bacterial alginate suspension was dripped with a commercial Peristaltic Pump (PD 5006 flux: 3.3 mL* min^−1^, Heidolph^®^, Schwabach, DE) with a Heidolph orbital agitator at 120 rpm for 30 min, using a syringe of 10 c.c. in a solution of CaCl_2_ at 3.2% with constant agitation; then, the forming macrobeads were washed with sterile saline solution at 0.85% (Gonzalez and Bashan 2000).

#### 2.4.2 Drying

Macrobeads were placed to dry on filter paper in sterile trays at 35°C, 40% of ventilation for 24 h, using a drying oven (Memmert UF55, Schwabach, Germany) and then maintained at room temperature in serological bottle containers until use.

#### 2.4.3 Selection of polymer-bacterium ratio

To select the polymer-bacterium ratio, a completely randomized design was used with a factorial 2*2 arrangement. The factors were polymer-bacterium ratio (80:20 and 70:30 vol / vol) and microorganisms (XT13 and XT14) immobilized as described in section 2.4.1. The response variable for the best polymer-bacterium ratio was measured by weight loss and cell variability of macrospbeads in soil.

#### 2.4.4 Visualization by scanning electron microscopy (SEM)

To visualize *Bacillus* sp. XT13 and *Bacillus megaterium* XT14 strains within the polymeric matrix, scanning electron microscopy (SEM) was performed. Samples (transversally cut macrobeads) were fixed with glutaraldehyde at 5% in sodium cacodylate buffer 0.1 M (pH 7.2). Subsequently, the samples were dried with CO_2_ to a critical point with a dryer (Samdri-PVT-3B, Tousimis, Rockville, MD, USA). The dry samples were placed on a sample holder and covered with 30 nm gold-palladium ratio of 60:40 in a sputter-coater engine (Vacuum Desk II, Denton, Scotia, NY). Visualization was performed in a scanning electron microscope (S-3000N, Hitachi, Tokyo, JP) at 15 kV, using an angle of 45° (image angle at the electron beam). The photographs were processed with software (Quartz PCI 5.5, Quartz Imaging, Vancouver, BC, CAN) (de-Bashan et al. 2016).

#### 2.4.5 Degradation of alginate macrobeads and cell viability

Macrobeads (0.2 g) were weighed and placed on very fine 5 × 5 cm (10-20 mm) nylon mesh, and 2.5 cm were placed under the soil surface in plastic pots (90 × 80 × 60 mm) using the procedure described by Bashan et al. (2002). Non-sterile soil collected in Cesar Department, Colombia was used for the experiment, which was also commercially used for guinea grass sowing, and whose physical-chemical characteristics were the following: pH (7.15), organic matter (3.02%), effective coefficient for cationic exchange (15.23 cmol / kg), P (215.93 ppm), K (0.63 cmol / kg), S (15.08 ppm), Ca (13.37 cmol / kg), Mg (1.8 cmol / kg), Na (0.05 cmol / kg), Cu (1.7 ppm), Fe (38.5 ppm), Mn (2.2 ppm) with analysis performed by the Analytical Chemistry Laboratory at Agrosavia.

The experiment was incubated in a phytotron at a temperature of 29 ± 2 °C, 60% humidity and a photoperiod of 16:8 h of light:darkness. Sterile distilled water addition for each one of the plastic pots was 2 mL/week. The experimental design consisted of four treatments T1: XT13 (80:20), T2: XT13 (70:30), T3: XT14 (80:20), T4: (70:30) each of them performed with three replicates and repeated twice. Sampling days were (0), (1), (5), (10) and (15). The response variables of this experiment were (a) macrobead weight of each treatment and (b) cell viability of the macrobeads of each treatment.

#### 2.4.6 Determination of cellular concentration

The serial dilution technique was used to determine number of viable cells (total CFU/ g of macrobeads) in TSA medium starting from immobilized spores (see section 2.3.1.2); then, 0.2 g of macrobeads were dissolved in 20 mL of sodium citrate (4% p/vol) and agitated at 300 rpm for 45 min. Subsequently, it was centrifuged at 4.637 × g for 10 min. Immediately, the spore pellet was resuspended in 1 mL of NaCl at 0.85%; aliquots of 100 μl were taken adding them to 900 μl of NaCl at 0.85%, performing subsequent dilutions up to 10^−8^. Then, from the 10^−8^ dilution, aliquots of 100 μl were plated in Petri dishes with trypticase soy agar (TSA, Difco Laboratories, Detroit, MI, USA.) culture medium, which were incubated at 30 °C for 24 h for subsequent counting.

### 2.5 Experiment in Guinea grass (*Megathyrsus maximus*) plants in stress condition by drought

The experiments were performed in the Centro de Investigación de Motilonia de Agrosavia located in Departamento del Cesar, Colombia in a greenhouse with semi-controlled conditions. Based on the experiments previously described, the formulation of the chosen prototypes for this trial were *Bacillus* sp. XT13 in Na-Alg at 3.2% and a polymer:bacterium ratio of 70:30 while *B. megaterium* XT14 was used in a polymer:bacterium ratio of 80:20 with the same Na-Alg percentage as that for strain XT13. Polyethylene 2-kg bags were used and filled with 1.5 kg of soil with the same chemical characteristics mentioned previously. The *M. maximus* var. Jacq guinea grass seeds were sterilized in surface with sodium hypochlorite at 3% for three min and rinsed three times in sterile water. Inoculation was performed at the moment of sowing. Treatments 3, 4 and 5 were inoculated with 5 mL of each bacterium at a concentration of 10^8^ CFU* mL^−1^. Treatments 6,7 and 8 were applied 0.2 g of sodium alginate mixed with 0.5 g of guinea grass seeds. Subsequently, water irrigation of 188 mL was applied per plant every day until they were established (24 days). Then, to induce them to a moderate drought stress, 94 mL of water (that is, 50 ± 5% in field capacity weight) were applied every four days until day 99. Irrigation was stopped immediately for six days to subject plants to a severe stress (without water). In total, the experiment lasted 105 days. Then, the destructive sampling was performed, and the foliar tissue of each one of the treatments was frozen in liquid nitrogen; samples were prepared for determining enzymatic activity of ascorbate peroxidase (APX) and proline. Additionally, values of the morphometric variables of dry weight of the aerial biomass part and root were taken. Finally, protein content, digestibility and neutral digestible fiber (NDF) were taken following the Near Infrared Spectroscopy (NIRS) method (Hopkins et al. 2019).

A completely randomized design was used with eight treatments and eight replicates per treatment. The treatments were (1) irrigated abiotic control; (2) abiotic control with drought stress; (3) XT14 nonpolymeric formulation with drought stress; (4) XT13 nonpolymeric formulation with drought stress; (5) XT14 plus XT13 without polymeric formulation with drought stress; (6) XT13 and polymeric formulation with drought stress; (7) XT14 and polymeric formulation with drought stress; and (8) XT14 plus XT13 and polymeric formulation with drought stress. In the treatments with the application of XT13 plus XT14, each strain was produced and immobilized separately and mixed in a ratio of 1:1.

#### 2.5.1 Enzymatic activity of peroxidase ascorbate (APX) and proline content

Plant material (120 mg) was macerated with liquid nitrogen; the enzymatic activity of ascorbate peroxide was determined by the method of Nakano and Asada (1981) and proline concentration by the method of Bates et al. (1973).

### 2.6 Statistical analysis

The data for the statistical analysis of the alginate degradation capacity, alginolytic activity, determination of the APX enzymatic activity, proline, morphometric variables of the aerial biomass part and protein content in root dry weight, digestibility and NDF were subjected to a statistical analysis with SAS statistics (Version 9.4) by means of one-way analysis of variance (ANOVA, *P* < 0.05) followed by the HSD Tukey’s test (post-hoc analysis).

Meanwhile, the two response variables used in the degradation test and cell viability of the macrobeads in soil were analyzed as follows: (a) Macrobead weight was performed with an analysis of variance (ANOVA, *P* < 0.05) followed by HSD Tukey’s test (post-hoc analysis) with SAS (version 9.4); (b) Cell viability of the macrobeads was performed by estimating the rate of change of the response variable in function of time for each treatment using a simple lineal regression analysis for each one of the repetitions by means of equation (2):

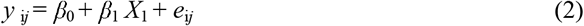

Where: *y*_i*j*_ : corresponds to the variable observed; *β*_0_ : is the interceptor for the regression; *β*_1_ : is the regression parameter for the independent variable (time); *X*_1_:is the independent variable (time); *e*_i*j*_ : is the error term in the regression.

Subsequently, the effect of the treatments was compared on the regression parameters subjected to one-way analysis of variance (ANOVA) (equation 3); in the cases where significant differences were found, measurement separation was performed with Tukey’s multiple comparison test at a level of significance of *P* < 0.05. For both regression and variance analyses, residual normality and homocedasticity assumptions were measured by graphic methods. The analyses were performed with the program SAS statistics (Version 9.4) by the procedures PROC REG and PROC GLM for regression and variance analyses, respectively.

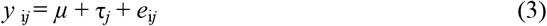

Where: *y*_i*j*_ corresponds to the estimated regression parameters; *μ*: corresponds to the general measurement; τ_*j*_ : is the treatments effect factor; *e*_i*j*_ : is the error term.

## 3. Results

### 3.1 Formulation of polymeric inoculants of *Bacillus* sp and *Bacillus megaterium*

Prototypes were formulated with a concentration of 3.2% of sodium alginate; macrobeads from 1 to 2 mm in diameter (Fig 1a) were obtained, in which *Bacillus* sp. XT13 and *B. megaterium* XT14 were immobilized with cellular concentrations higher than 10^9^ CFU mL^−1^ in 0.2 g of dry macrobeads alginate. Likewise, within the polymeric matrix, the bacterial homogenization was observed by scanning electron microscopy (SEM). (Fig 1b,1c).

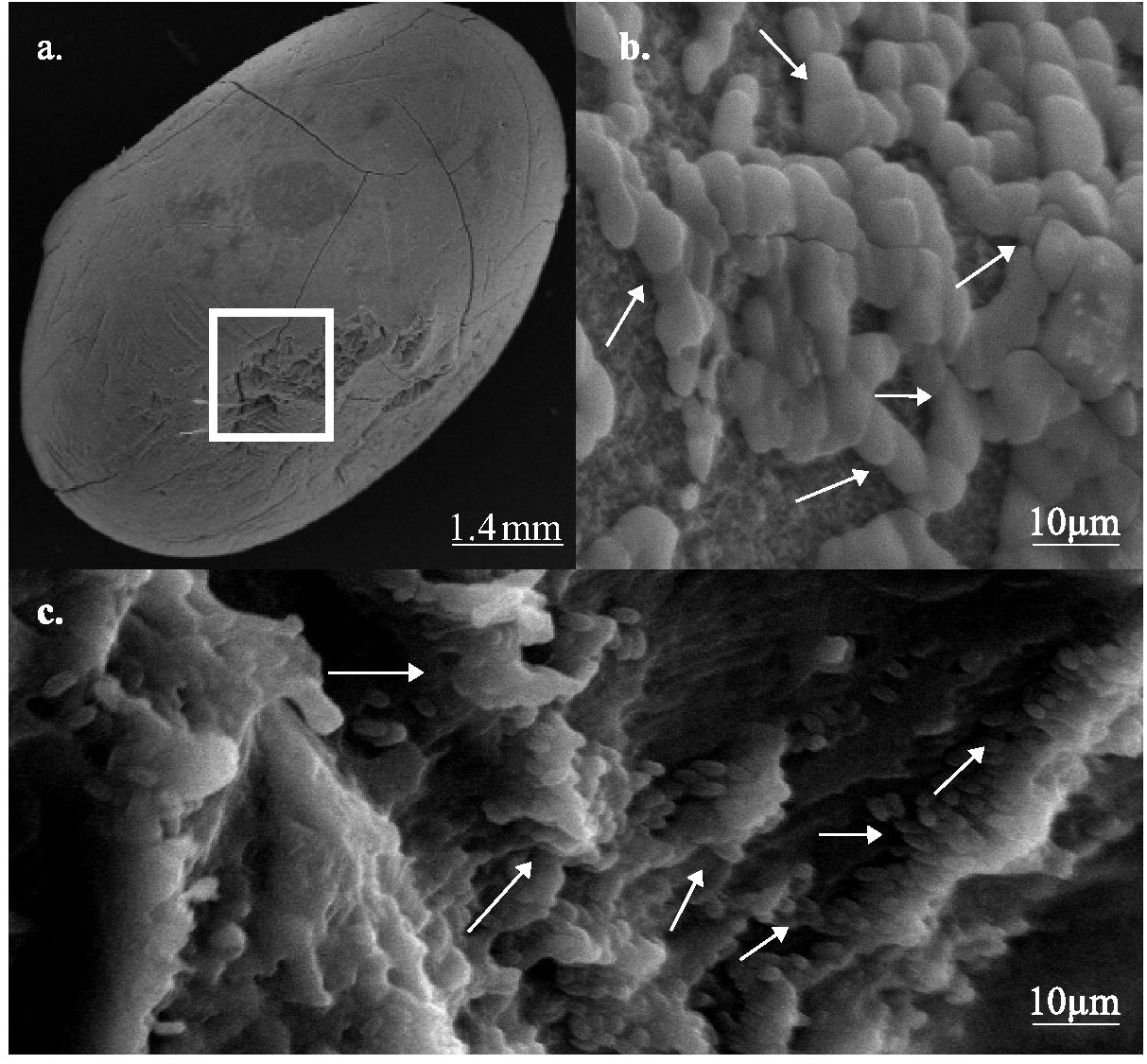

### 3.2 Degradation capability and alginate consumption

None of the strains was capable of growing in M.A. culture medium without alginate (Fig 2). In contrast, the results in M.A medium supplemented with alginate as the only carbon source showed that *Bacillus* sp. XT13 significantly increased the optical bacterial density (*P* < 0.05).

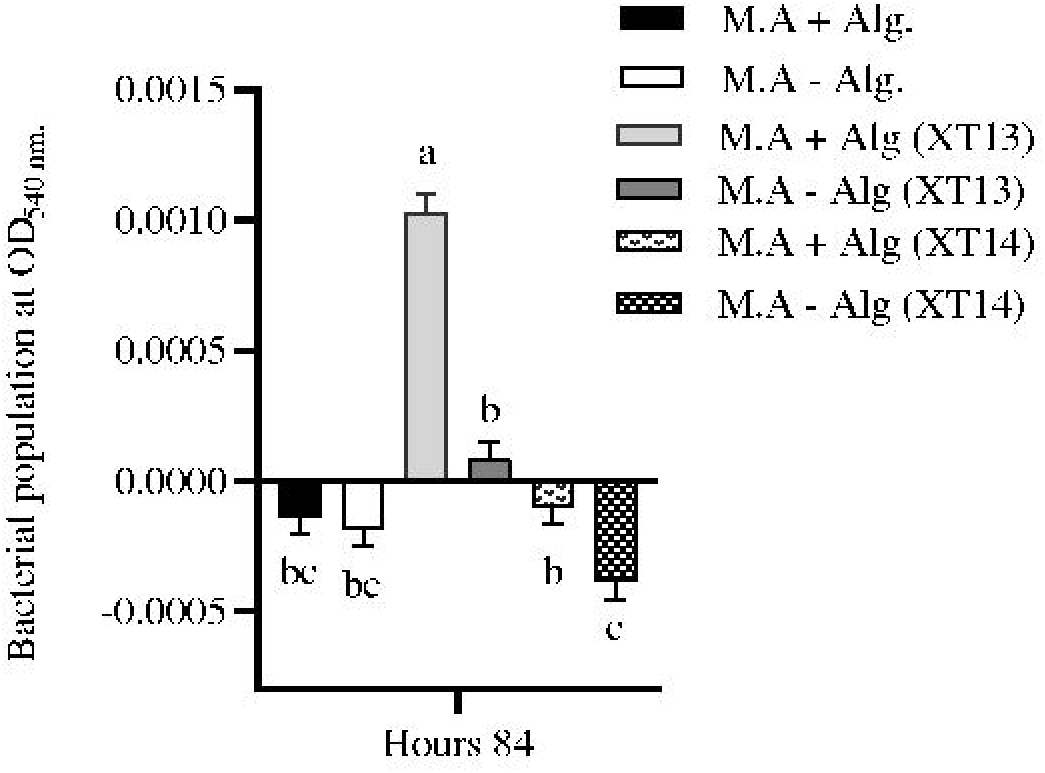

In the assay of alginate degradation, a decrease of 9.4% viscosity was observed in the control group without bacteria at 24 h. At the other sampling times, viscosity was observed constant (49.17 ± 1 cP; Fig. 3a, b). Additionally, the alginate + *Bacillus* sp. XT13 (80:20) and alginate + *Bacillus* sp. XT13 (70:30) treatments showed a progressive decrease of 5.90 % and 3.07 % of alginate viscosity, respectively (Fig.3a). For *Bacillus megaterium*. XT14 strain, no evidence was observed of viscosity decrease in any of the two ratios (Fig.3b).

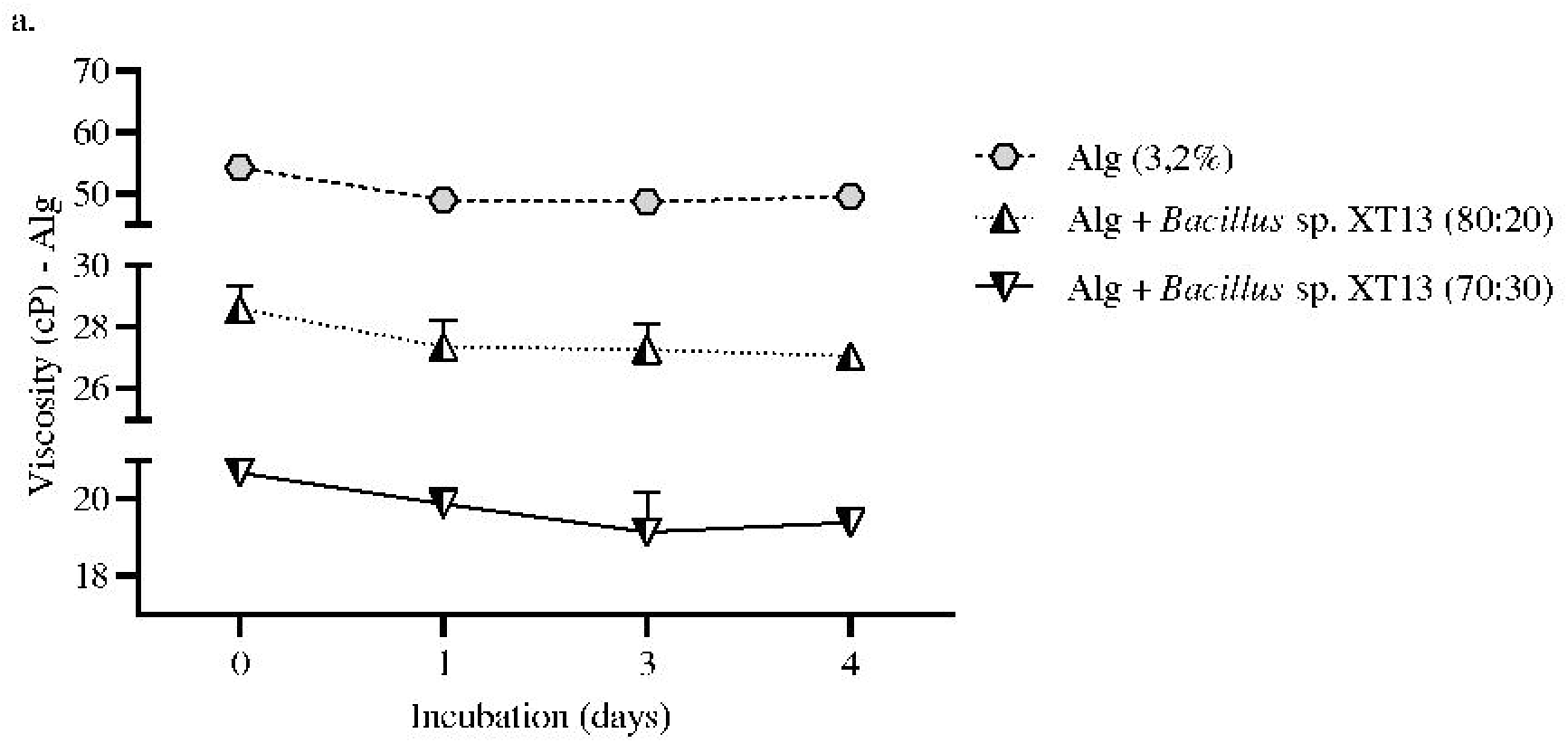
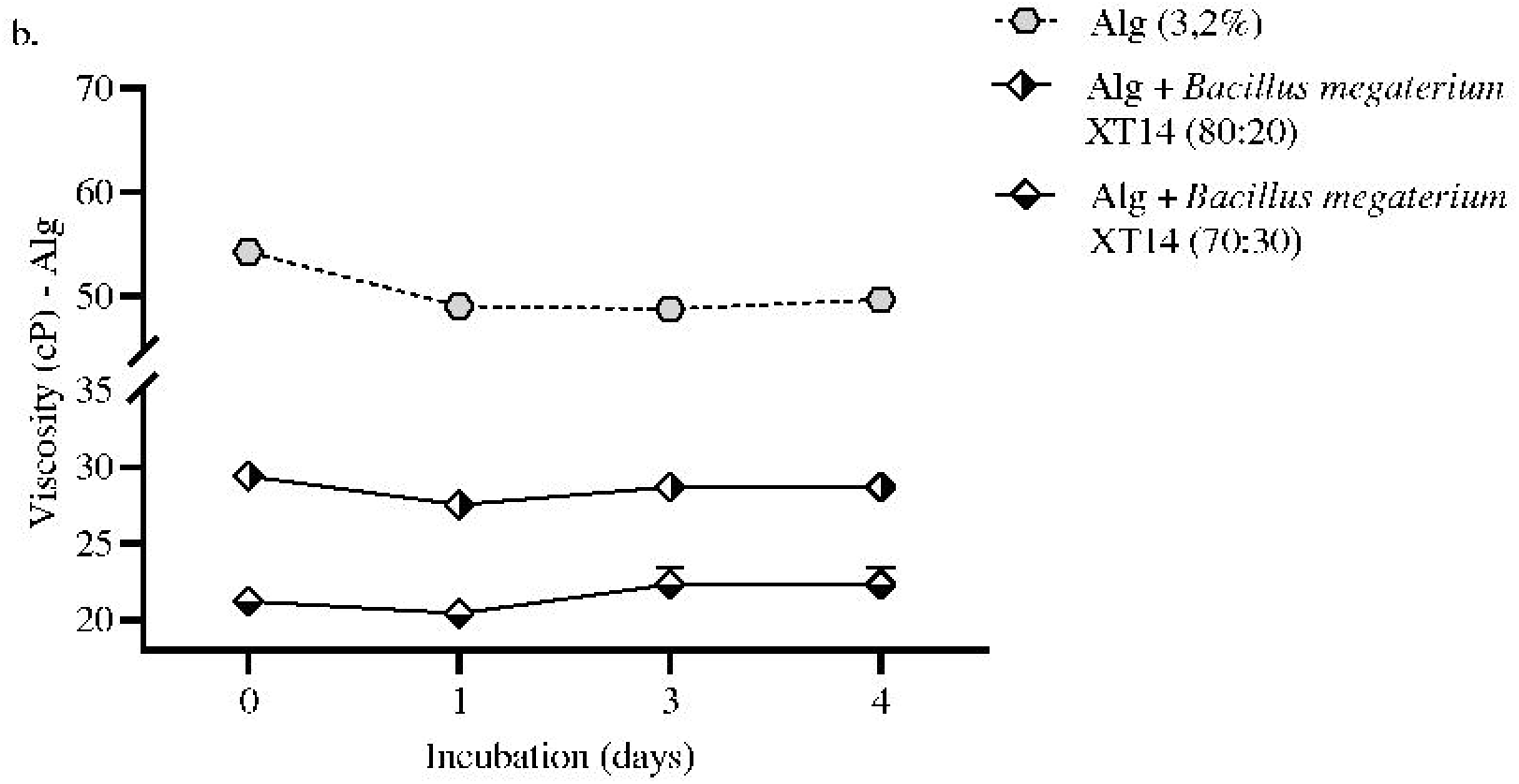

### 3.3 Degradation of alginate macrobeads and cell viability

At the end of the test (15 days), the results obtained showed significant (*P* <0.05) differences in the two variables assessed: weight loss (g) of the macrobeads and cell viability of the macrobeads in soil, indicating tehir possible degradation; the most rapid and evident degradation of the macrosbeads in soil was in the proportion polymer:bacterium 70:30 with *Bacillus* sp XT13 with a weight loss of 0.06 g and a decrease in cell viability of 20.5% equivalent to 2.513 UFC mL^−1^, and whose slope value (m) was m: −0.142. The proportion 80:20 followed with *Bacillus megaterium* XT14, showing a weight loss of the macrobead of 0.03 g and a decrease of cell viability of 16.74% equivalent to 1880 UFC mL^−1^ and whose slope value (m) was m: −0,105 (Fig. 4 a, b).

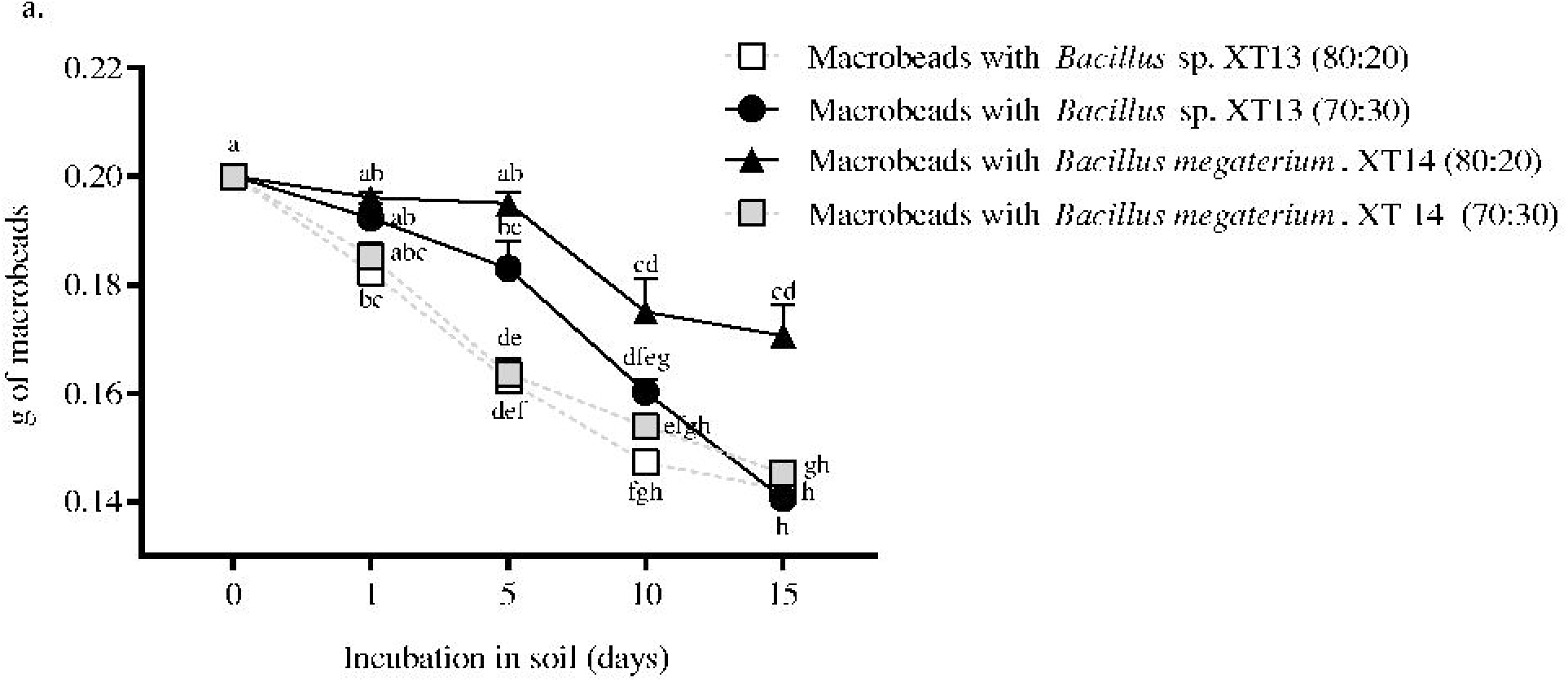
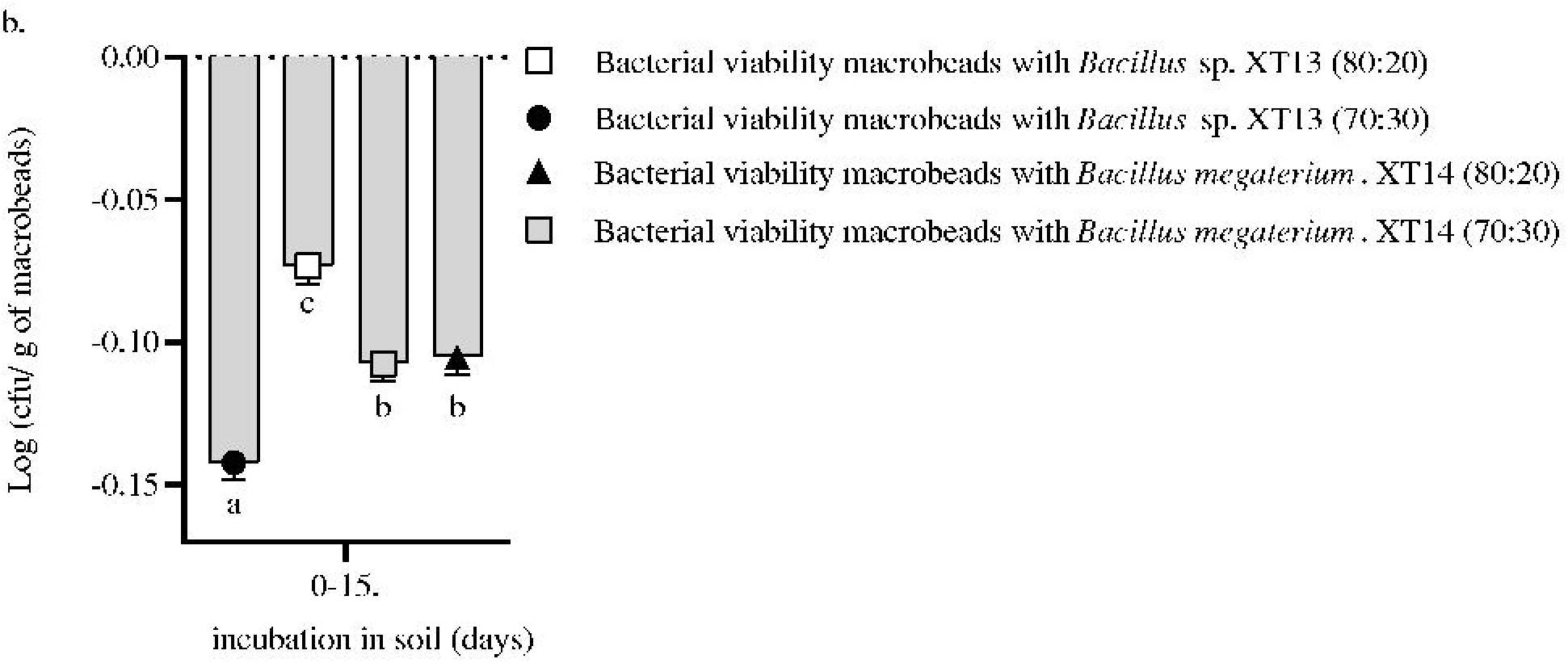

## 4. Drought tolerance induced in Guinea grass by *Bacillus* sp. and *Bacillus megaterium* immobilized in alginate macrobeads

### 4.1 Effect on dry biomass production

The treatment with the greatest significant effect on dry biomass of the aerial part was evident in the bacterial mix (*Bacillus* sp. XT13 + *Bacillus megaterium* XT14) immobilized with an increase of 1.8 times compared with the control treatment under drought. Subsequently, the treatments *Bacillus megaterium* XT14 immobilized and *Bacillus* sp. XT13 immobilized increased 1.6 and 1.5 times this response variable, respectively. Likewise, the mix of immobilized microorganisms (*Bacillus* sp. XT13 + *Bacillus megaterium* XT14) increased 3.3 times the root dry biomass, followed by *Bacillus megaterium* XT14 immobilized (3.1 times) and *Bacillus* sp. XT13 immobilized (2.6 times) compared with the non-innoculated drought treatment (Table 1).

### 4.2 Effect on nutritional quality of Guinea grass (*Megathyrsus maximus*)

For the three response variables related with nutritional quality (crude protein, neutral digestible fiber and digestibility), a similar tendency was observed in all the treatments. The results showed that crude protein and digestibility of guinea grass increased 20% and 3% when the microorganisms immobilized in macrobeads were applied, compared with the non-immobilized strains; similarly, in neutral digestible fiber (NDF), a decrease of 4% was found in this same condition (Fig. 5a and 5b). The treatment with the greatest beneficial effect (*P* < 0.05) was the mix (*Bacillus* sp. XT13 + *Bacillus megaterium* XT14) immobilized with 9.58%, 64.93% and 56.8% in crude protein, NDF and digestibility where the non-inoculated treatment under drought obtained 3.15%, 69.02% and 51.32%, respectively.

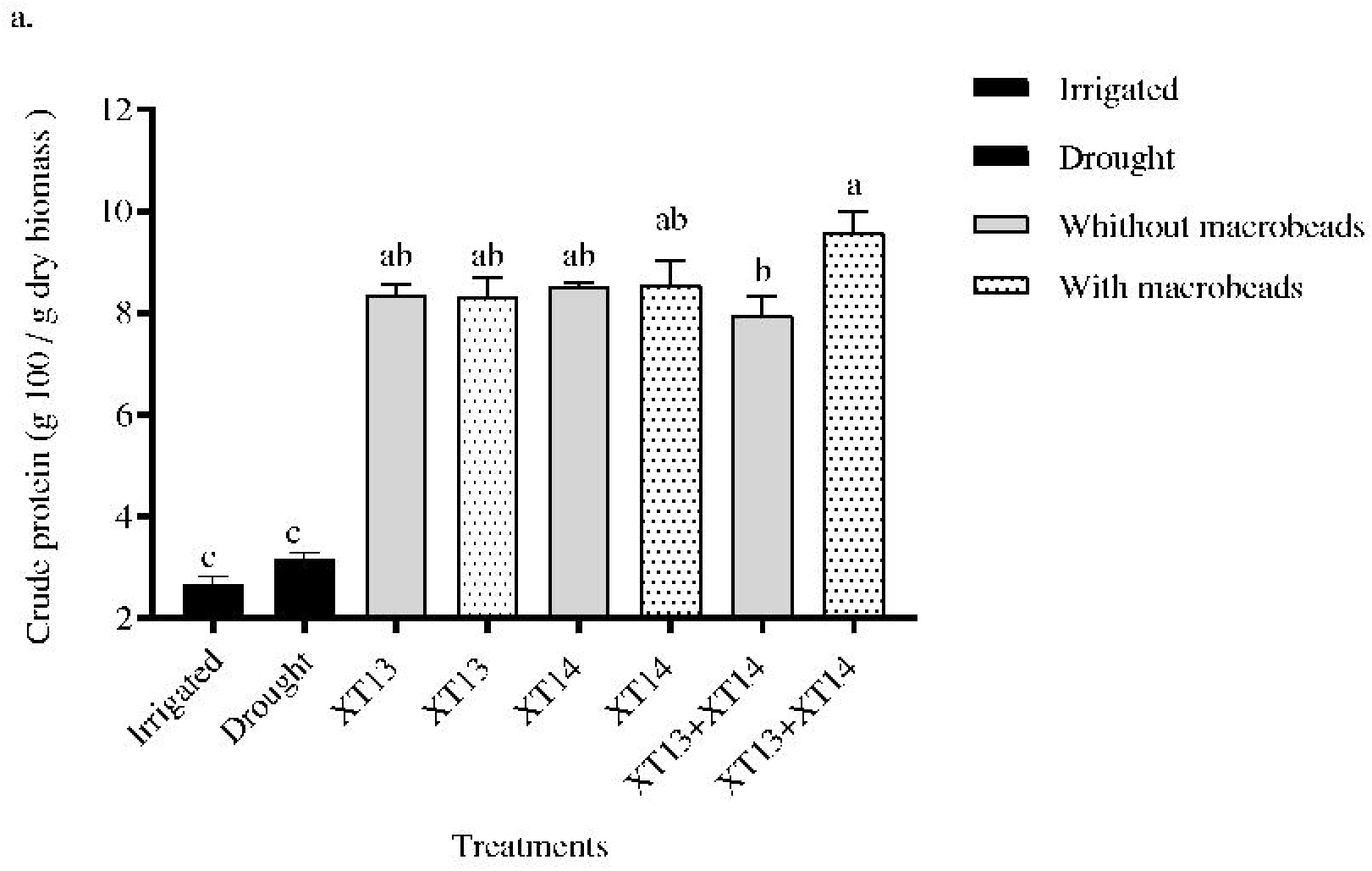
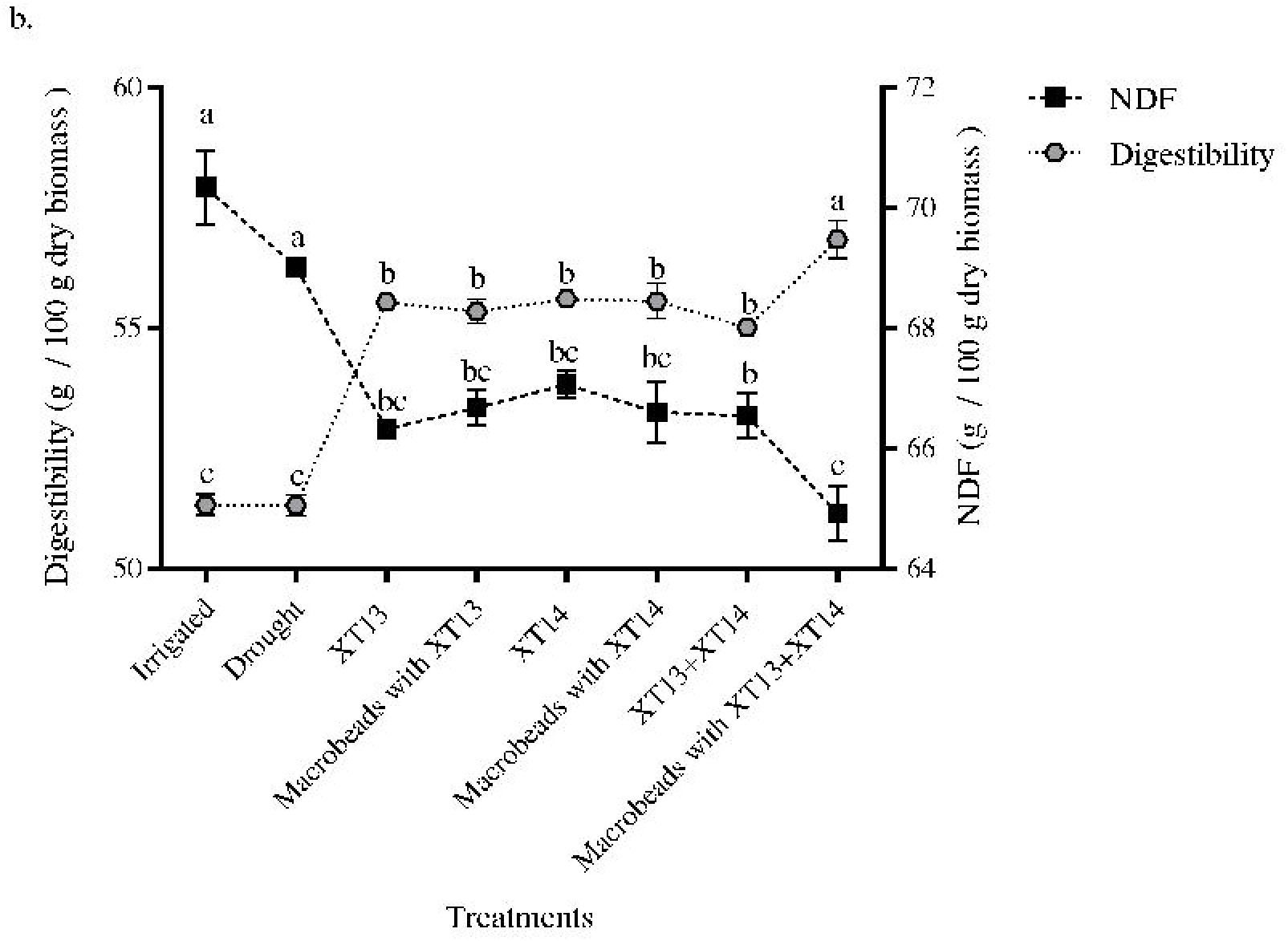

### 4.3 Effect of the response on proline accumulation and antioxidant activity

A significant effect (*P* < 0.05) in proline accumulation was observed in three treatments under immobilization in macrobeads: *B. megaterium* XT14, *Bacillus* sp. XT13 and *Bacillus* sp. XT13 + *B.megaterium* XT14 with 89.77, 84.58 and 46.73 μg proline/ mg of plant tissue, respectively, compared with the non-inoculated control treatment under drought with 27.16 μg proline/ mg plant tissue (Fig. 6a).

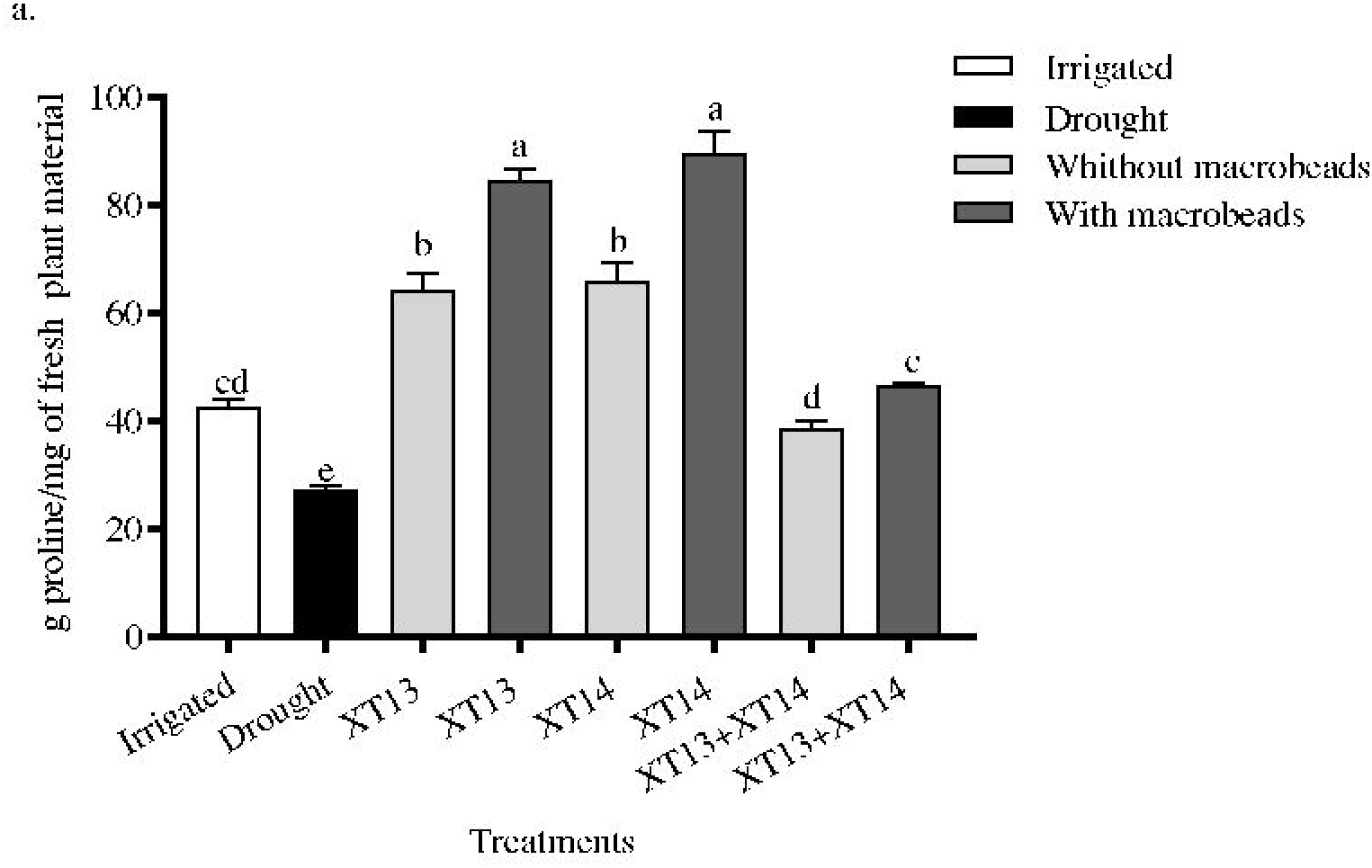
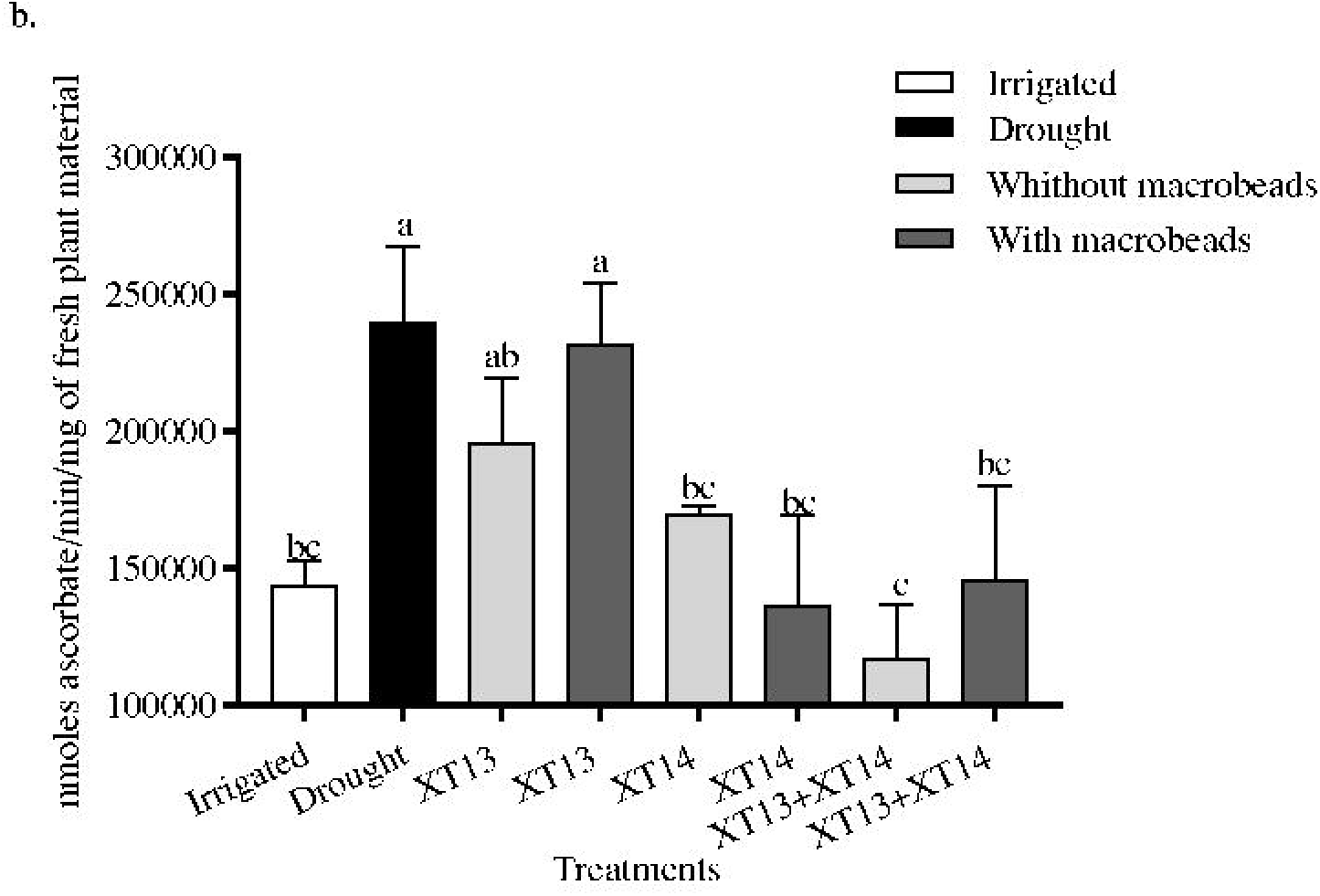

Moreover, statistical differences (*P* < 0.05) were measured among several treatments while studying the antioxidant response of APX in guinea grass. The application of the non-immobilized bacterial mix produced 117.5 nmoles of ascorbate/min/mg of plant tissue compared to the control under drought with 239.69 nmoles ascorbate/min/mg of plant tissue. The two treatments that followed with significant influence were *Bacillus megaterium* XT14 immobilized and the immobilized microorganism mix with 136.37 and 146.07 nmoles ascorbate/min/mg of plant tissue (Fig. 6b).

## Discussion

A promising alternative to counteratack the negative effects caused by drought in Guinea grass forage quality and quantity is the development of inoculants under polymeric formulation. Thus, the main objectives of this study were (1) evaluate and determine the necessary conditions to immobilize the strains of interest in dry macro-polymeric matrix and (2) investigate the influence of bacterium immobilization in macrobeads on increasing tolerance to drought stress in Guinea grass. The strains used in this research, *Bacillus* sp. XT13 and *Bacillus megaterium* XT14, were isolated and characterized as plant growth-promoting bacteria (PGPB) with xerotolerant capacity in previous research in the Agricultural Microbiology Laboratory of Agrosavia (Moreno et al. 2015). Additionally, Abril et al. (2017) demonstrated that under greenhouse conditions, XT13 and XT14 were capable of inducing tolerance to drought in Guinea grass improving its growth. Besides microorganisms, polymer has a significant role in the immobilization process for dry macro-polymeric matrix production. In this study, we used sodium alginate and performed experiments under conditions *in vitro* to evaluate and determine the necessary conditions to immobilize the strains of interest in dry macro-polymeric inoculant. We hypothesized that if the strain is capable of consuming and degrading alginate, the polymeric matrix biodegradability will be greater, which will be observed both with the progressive decrease in weight and cell viability of the macrobeads. To our knowledge, very few references are available with results related to aspects of PGPB and alginate interaction, which is a complex co-polymer of α-L-guluronate (G) and β-D-mannuronate (M) (Tang et al. 2009). Thus, the first step was to demonstrate that strains, XT13 and XT14 were capable of consuming and degrading alginate (see 2.3).

To evidence alginate consumption, optical density (DO_540 nm_) was measured for 84 h as a bacterial growth indicator, using a minimum medium (M.A) supplemented with alginate as the only carbon source. The only strain that showed an increase in DO_600 nm_ was *Bacillus* sp. XT13. In contrast, no other strain was capable of growing in the M.A medium without alginate, which suggests that *Bacillus* sp. XT13 is capable of consuming alginate (Fig.2). As an indirect measure to infer if the microorganisms degraded alginate, polymer viscosity was assessed through time in the presence of each bacteria in a concentration of alginate (3.2%), which was selected in previous studies (data not shown). Likewise, two polymer:bacterium (70:30 and 80:20) ratios were used in this experiment as reported by (Abd El-Fattah et al. 2013; Wu et al. 2011). The control without bacteria only showed a non-significant decrease of alginate viscosity at 24 h, likely caused by shear forcing generated at 100 rpm. During the other sampling times, the viscosity in this treatment was constant. In the treatments with bacteria, the only strain that decreased (*P* < 0.05) alginate viscosity was *Bacillus* sp. XT13 in both polymer: bacterium ratios, suggesting that this strain have alginolytic activity (Fig 3a and 3b). Alginates can be disassembled by alginate lyases to monosaccharides or alginate oligosaccharides, and the latter exhibits many fascinating bioactivities (Wang et al. 2012). According to Michel et al. (2006), the metabolic capacity to degrade and be used as carbon source in alginate has been little studied and observed only in bacterial genera, such as *Pseudomonas* sp. and *Bacillus* sp. These results indicated that *Bacillus* sp. XT13 was capable of degrading and consuming alginate.

Bashan et al. (2014) stated that a key objective in microorganism immobilization was to be able to control the release of soil bacteria to allow plant inoculation for a period of time. The parameters that influenced directy in such objective were (i) polymer concentration and (ii) polymer: bacterium ratio (Bashan 1998). As mentioned previously, we used the concentration of 3.2% alginate, obtaining macrobeads from 1 to 3 mm besides the polymer: bacterium 70:30 and 80:20 ratios in each one of the bacterial strains. The selection of the most beneficial ratio for each strain was determined by macrobead cell release. (He et al. 2016) and Bashan et al. (2014) reported that a gradual release of PGPB, such as *Pseudomonas putida* Rs-198 and *Azospirillum brasilence* of the polymer matrix was given by the property of alginate of being degraded because it is a polysaccharide that can be used as carbon source by the strains previously mentioned and native soil microbial communities. According to Liakos et al. (2014), alginate degradation occurrs possibly through the breaking of covalently linked (1–4) glycoside bonds of sodium alginate composed of unbranched chains β-dmannuronate (M) and α-l-guluronate (G) residues. For this reason, we measured weight of the macrobeads and cell viability (UFC mL^−1^) through time selecting the ratio that could produce the greatest cell release.

As indicated in Bashan (2016), slow or fast release of PGPB/PGPR immobilized in alginate macrosbeads is related to polymer degradation in soil. Thus we observed a faster release of *Bacillus* sp. XT13 in the polymer:bacterium 70:30 ratio whose weight loss of the macrobeads was 0.06 g (equivalent to 2.382 UFC mL^−1^) and 0.058 g (equivalent to 1.657 UFC mL^−1^) in the ratio 80:20 during the 15-day assay, so we selected 70:30 for *Bacillus* sp. XT13 to perform the immobilization process. With respect to *Bacillus megaterium* XT14, the fastest release was observed with the ratio 80:20 whose weight loss was 0.03 g (equivalent to 1.880 UFC mL^−1^) and 0.05 g (equivalent to 1.550 UFC mL^−1^) in the ratio 70:30. Thus, we selected the ratio 80:20 to perform the immobilization process of *Bacillus megaterium* XT14 strain. The biodegradation rate of the polymer matrix was directly related to its composition, preparation method and biological activity of the microorganisms (Bashan et al. 2016; Cortes-Patino and Bonilla 2015). With respect to the biological activity, the results previously described and those reported by Vogelsang and Østgaard (1996), the alginate exposed to non-sterile systems, such as wastewater and soil, can be degraded by microorganisms. We cannot exclude the possibility that native microbial communities in soil could have an influence on alginate biodegradability in this study. Interstingly, between the two microorganisms, we confirmed that treatments with *Bacillus* sp. XT13 showed greater weight loss of the microbead and a high cell release of the polymer matrix, which indicated that the greatest microbead biodegradation was obtained with this strain. Curiosly, XT13 showed the ability of consuming and degrading alginate under *in vitro* conditions. Based on these findings, we confirm the first hypothesis set out in this study.

Subsequently, we carried out a greenhouse experiment to study the effect of bacterial immobilization in macrobeads on drought tolerance induced in Guinea grass. We evaluated Guinea grass response to individual inoculation and co-inoculation of *Bacillus* sp. XT13 and *Bacillus megaterium* XT14 strains under drought conditions. Initially, the irrigation control and drought control treatments were compared to confirm the stress effects on the plant. We observed that drought stress did not affect biomass production and Guinea grass digestibility. On the contrary, dry root weight decreased (*P* < 0.05) in drought. In this study, we also report the impact of drought stress and PGPB inoculation on compatible osmolytes and activities of enzymatic antioxidants related to the redox status of Guinea grass. To accomplish this objective, proline accumulation and ascorbate peroxidase activity was assayed (Ghosh et al. 2018). In such treatments we evidenced that proline decreased and APX increased significantly. These results indicated that drought had a negative effect, but it was not significant on Guinea grass growth under the conditions of this study. According to Fang et al (2015), plants have stress resistance mechanisms at physiological and biochemical levels where *M. maximus* is not the exception (Borjas-Ventura et al. 2019). Despite this evidence, in the following comparisons we demonstrated the significant (*P* < 0.05) inoculation effect of *Bacillus* sp. XT13 and *Bacillus megaterium* XT14 with and without salt in dry macro-polymeric matrix.

Our experimental design also allowed us to corroborate the beneficial influence of the strains on Guinea grass growth in dry conditions. For this purpose, we compared the inoculated plants without microbead immovilization against those with bacterial application under stress conditions. We observed the strains increased biomass production significantly (*P* < 0.05) in the aerial part, root biomass, proline and protein in 75, 174, 106 and 162%, respectively. In addition, digestibility improved 7.6% and decreased NDF percentage to 2.6%. Furthermore, while the ascorbate peroxidase antioxidant activity recorded the highest levels under water stress, inoculation with XT13 and XT14 caused a significant reduction in APX activity (32.4%). These results indicated that the strains under study were capable of promoting Guinea grass growth in drought conditions as reported in Abril et al. (2017) and Moreno et al. (2015). Similar observations have been described in the few investigations reported on the use of PGPR for Guinea grass without hydric stress by drought. For example, Mishra et al. (2008) found that *Azospirillum brasilense* and an arbuscular microrrhiza fungus consortium increased germination percentage (> 90%), crude protein production (> 7%) and improved NDF (> 30%). The relevance of these results is that they were obtained under drought stress using response variables, such as antioxidant enzymatic activity (APX), osmoprotector osmolite accumulation (proline), nutritional quality parameter (NDF, protein digestibility) and biomass production. Proline accumulation was assessed in this study because it plays many roles, such as preventing water exosmosis by decreasing cell water potential (Reddy et al. 2015), acting as a source of energy (Szabados and Savouré 2010), maintaning cytosolic pH and intracellular redox potential (Ben Rejeb et al. 2014), stabilizing protein structure through chaperon activity (Liang et al. 2014) and preventing oxidative damage under abiotic stress conditions (Ghosh et al. 2018). Likewise, APX is found in the chloroplasts, mitochondria, peroxisoma and cytosol detoxifying the hydrogen peroxide generated under drought conditions (Miller et al. 2010). Additionally, crude protein, NDF and digestibility are feed parameters related with degradability and nutritional contribution of livestock forage (Delevatti et al. 2019). Finally, the importance of biomass production lies in being the main food source for livestock. Pertinent reports have suggested that PGPB can act mainly in two ways on plants under drought conditions. The first PGPB role consists on modulating antioxidant machinery, in addition to other components in plant systems leading to altered metabolic fluxes and inducing systemic tolerance. The second role is that growth promotion activities, such as EPS production and indole acetic acid synthesis, metabolic capabilities shown by XT13 and XT14 (Abril et al. 2017), influence increased protection against desecation, microbial aggregation and stimulate root elongation (Naseem et al. 2018). Therefore, growth promotion confirmed in this experiment allowed us to infer that *Bacillus* sp. XT13 and *Bacillus megaterium* XT14 induced drought stress tolerance on Guinea grass.

Subsequentely, we investigated the effect of the immobilization process on the capacity of XT13 and XT14 to induce hydric stress tolerance. We hypothesized that macrobead immobilization had a positive influence in the capacity of XT13 and XT14 to induce hydric stress tolerance. To test this idea, we compared the inoculated plants with immobilized bacteria in dry macro-polymeric matrix against those treatments where bacterial application was performed without immobilization. For dry measured biomass, we confirmed increases of 7.32 and 25.3% in the aerial part and root when the immobilized mix of *Bacillus* sp. XT13 and *Bacillus megaterium* XT14 was applied. On the other hand, a greater beneficial effect of the strains was observed when they were immobilized under the nutritional quality parameters performed because digestibilitiy increased 3.32%, NDF decreased 2.43% and protein increased 20.3% when co-inoculation (XT13+XT14) was applied. With respect to proline osmoprotectant osmolyte, we noted a significant increase in accumulation of 31.3, 36.07 and 21.06% when applied to each treatment *Bacillus* sp. XT13, *Bacillus megaterium* XT14 and *Bacillus* sp. XT13 + *Bacillus megaterium* XT14, respectively. Additionally, a reduction in reactive oxygen species (ROS) synthesis was evident by the action of the antioxidant enzyme APX in plants under drought conditions that were immobilized with PGPB in dry macro-polymeric inoculant. The APX response was significant (*P* < 0.05) with the immobilized *Bacillus megaterium* XT14 and the immobilized mix *Bacillus* sp. XT13 + *Bacillus megaterium* XT14 treatments with a decrease of 19.6 % and 24.2 % compared to that obtained with the non-immobilized organisms. These results corroborated those reported by Ghosh et al. (2018) recording the highest level in all the redox (ROS molecules and antioxidant enzymes) under hydric stress; inoculation with *Pseudomona putida* GAP-P45 decreased ROS accumulation and all antioxidant enzyme activities including (APX) in seedlings of *A. thaliana* significantly (*P* < 0.05) in the majority of the analysis points in time condition of water deficit. Based on these findings, we inferred that Guinea grass showed better growth under drought when PGPB were immobilized in macrobeads. In this sense, and considering the results mentioned above, our study provides evidence that inmobilizing PGPB in macrobeads influences stress tolerance beneficially, which confirms our hypothesis.

An explanation of the results found might be related with the protection and gradual release of PGPB provided by the polymeric matrix. These two effects decreased the negative influence of extreme environmental factors and native microbial competence on XT13 and XT14, possibly extending tolerance in Guinea grass longer. Recent evidence has demonstrated that the use of integrating super absorbent polymers (SAP) with PGPB in liquid cultivations is a viable stratey to mitigate the effects of drought in wheat and cucumber (Li et al. 2019). In addition, other studies in China have demonstrated that poly-γ-glutamic acid (γ-PGA) synthesis by the two *Bacillus subtilis* and their application increased resistance to drought in maize improving growth of microbial populations *Bacillus, Pseudomonas, Burkholderia, Talaromyces* and *Rhizopus* in soil (Yin et al. 2018). In these two studies, the authors used a liquid formulation with polymeric additives, which are formulations of the first generation and different to that demonstrated in this research. In this study we used a second generation encapsulated polymeric formulation (Bashan et al. 2014) where *Bacillus* sp. XT13 and *Bacillus megaterium* XT14 strains were immobilized in the alginate matrix. Additionally, in this study the dry macro-polymeric matrix treatment without bacteria was not evaluated because alginate does not generate any effect on the plant as shown in tomato, wheat and mesquite (Bashan et al. 2002; Gonzalez et al. 2018).

Finally, based on all the comparisons described, we deduced that co-inoculation of *Bacillus* sp. XT13 and *Bacillus megaterium* XT14 where each strain was immobilized in macrobeads was the treatment with the greatest tolerance provided in general to Guinea grass under drought. Because *M. maximus* had a significant impact in silvopastoral systems in Colombia for the livestock sector, integrating the use of *Bacillus* sp. XT13 and *Bacillus megaterium* XT14, jointly with the polymeric formulation of alginate macrobeads, could be projected as a strategy to improve production of this grass facing the effect of lengthy drought due to climate change. To the best of our knowledge, this study is one of the first reports about the influence of synergistic use of PGPB xerotolerant and dry macro-polymeric formulation on Guinea grass production under drought stress.

To conclude, the results in this research have shown that (1) *Bacillus* sp. XT13 is capable of degrading and consuming alginate; (2) the polymer: bacteria ratio influences the gradual microbial release in the soil from macrobeads, which were 80:20 for strain XT14 and 70:30 for strain XT13 at a concentration of 3.2% alginate; (3) *Bacillus* sp. XT13 and *Bacillus megaterium* XT14 induced drought stress tolerance in Guinea grass; and (4) drought stress tolerance of Guinea grass significantly increases when the strains were inoculated and inmovilized using the dry macro-polymeric formulation. Extensive field studies are necessary to corroborate the response of Guinea grass under drought stress on *Bacillus* sp. XT13 and *Bacillus megaterium* XT14 inoculation immobilized individually in alginate macrobeads. In addition, further research aimed at understanding how to perform a co-immobilization process of two or more microorganisms and clarify their interaction within the macrobead could represent an improvement in second generation bioferttilizers.

## Supporting information

Table 1. Drought and inoculation effect of Bacillus sp. XT13 and Bacillus megaterium XT14 non-immobilized and immobilized in alginate macrobeads on dr

## Acknowledgments

The authors thank Andrea Bernal and workteam of the Agencia Presidencial de Cooperación Internacional de Colombia (APC) and Agencia Mexicana de Cooperación Internacional para el Desarrollo (AMEXCID) for the funding provided for this study. Special thanks to Yoav Bashan, Emeritus Ph.D. in Science†, and Luz Estela González de Bashan for the training received to perform bacterial immobilization in macrobeads; to Ronnal Ortiz for statistical advising and Diana Fischer for translation and editorial services in English.

## Conflict of interests

The authors declare no conflict of interests

