## Supplementary material for "Enhancement of drought tolerance on guinea grass by dry alginate macrobeads as inoculant of *Bacillus* strains": Table 1. Drought and inoculation effect of Bacillus sp. XT13 and Bacillus megaterium XT14 non-immobilized and immobilized in alginate macrobeads on dr

Tables.

**Table 1.** Drought and inoculation effect of *Bacillus* sp. XT13 and *Bacillus megaterium* XT14 non-immobilized and immobilized in alginate macrobeads on dry biomass production of the aerial part and root dry weight of guinea grass. Drought experiment lasted 81 days (75 of moderate stress + 6 of severe stress).

| **Treatments** | **Shoot dry biomass (g/plant).** | **Root dry weight (g/plant).** |
|---|---|---|
| **Irrigated** | 3.31b* (±0.20) | 1.46 bc (±0.14) |
| **Drought** | 3.33b (±0.13) | 0.96c (±0.19) |
| **XT13 without macrobeads** | 5.75a (±0.42) | 2.53ab (±0.37) |
| **XT13 with macrobeads** | 5.31a (±0.35) | 2.50ab (±0.36) |
| **XT14 without macrobeads** | 6.26a (±0.18) | 2.78ab (±0.37) |
| **XT14 with macrobeads** | 5.53a (±0.34) | 3.03a (±0.31) |
| **XT13+XT14 without macrobeads** | 5.60a (±0.29) | 2.61ab (±0.41) |
| **XT13+XT14 with macrobeads** | 6.01a (±0.19) | 3.26a (±0.42) |

*According to the post hoc Tukey's test; values denoted by a different small letter differ significantly ($P < 0.05$) in one-way ANOVA (median ± standard error).
